# Analytical considerations for comparative transcriptomics of wild organisms

**DOI:** 10.1101/048652

**Authors:** Trevor J. Krabbenhoft, Thomas F. Turner

## Abstract

Comparative transcriptomics can now be conducted on organisms in natural settings, which has greatly enhanced understanding of genome-environment interactions. However, important data handling and quality control challenges remain, particularly when working with non-model species outside of a controlled laboratory environment. Here, we demonstrate the utility and potential pitfalls of comparative transcriptomics of wild organisms, with an example from three cyprinid fish species (Teleostei:Cypriniformes). We present computational solutions for processing, annotating and summarizing comparative transcriptome data for assessing genome-environment interactions across species. The resulting bioinformatics pipeline addresses the following points: (1) the potential importance of “essential genes”, (2) the influence of microbiomes and other exogenous DNA, (3) potentially novel, species-specific genes, and (4) genomic rearrangements (e.g., whole genome duplication). Quantitative consideration of these points contributes to a firmer foundation for future comparative work across distantly related taxa for a variety of sub-disciplines, including stress and immune response, community ecology, ecotoxicology, and climate change.

## INTRODUCTION

High throughput sequencing has dramatically accelerated the pace of genomic research. While once restricted to model species in laboratory settings, genomic methods are being widely applied to non-model species in nature, rapidly illuminating the black box of the genome and giving rise to the field of ecological genomics. Reduced sequencing costs have made it feasible to study transcriptomes of co-occurring species in a community ecology context (i.e., “community transcriptomics”), as well as comparative studies of transcriptome evolution across diverse clades (i.e., “comparative transcriptomics”). While genomic data from model species can be informative for the biology of related organisms, not all species are the same in terms of their ecology, genetics, and morphology. For example, research on the zebrafish *(Danio rerio,* family Cyprinidae) can be relevant for other members of this hyper-diverse clade, but cannot explain the tremendous ecological and morphological diversity in this clade, as studies of single species are insufficient for understanding dynamic interactions among species and their respective genomes in a macroevolutionary context. In order to understand the causes and consequences of those interactions, as well as the origin of evolutionary novelty (e.g., new genes), we must examine genomes across species that reflect that diversity.

Comparative transcriptomics of co-occurring species in the wild has enormous potential to advance our understanding of mechanisms underlying molecular adaptation, evolutionary diversification, and community ecology. In this context, several important questions arise. For example: What are the proximate and ultimate mechanisms underlying phylogenetic, ecological, and morphological divergence? How have ancestral genomes been molded by divergent natural selection and other evolutionary forces into myriad forms that exist today? How does genomic architecture constrain or promote diversification? How important are genome duplication events in adaptive radiations? What role do genomes play in underlying the ecological dynamics of community assembly (e.g., competition, abundance, spatial and temporal dynamics, physiological constraint, etc.)?

Comparative study of transcriptomes in the wild presents important bioinformatic challenges that must be addressed in order to produce quality assemblies and annotations. For example, the degree to which information from model-organism databases can be extended to related non-model species is unknown and sources of genetic material in a given sample need to be partitioned correctly (e.g., target species versus microbiome). In this paper, we illustrate a computational strategy for comparative analysis of transcriptomes from a hyper-diverse lineage (>2,000 species) of vertebrates, the family Cyprinidae. The family includes zebrafish *(Danio rerio)* (Briggs 2002; Dooley & Zon 2000; Lieschke & Currie 2007), a model species with a comprehensively-annotated genome (Howe *et al.* 2013). We evaluated the application and limitations of zebrafish resources for annotating transcriptomes of three evolutionarily related, but non-model species: *Cyprinella lutrensis* (red shiner), *Platygobio gracilis* (flathead chub) and *Cyprinus carpio* (common carp).

Transcriptomes of *Cyprinella lutrensis* and *Platygobio gracilis* have not been published to our knowledge, whereas genomic and transcriptomic data are available for *Cyprinus carpio* (Ji *et al.* 2012; Wang *et al.* 2012; Xu *et al.* 2014). *Cyprinella lutrensis* and *Platygobio gracilis* are diploid (2n = 50), while *Cyprinus carpio* is allotetraploid (2n = 100; Ohno *et al.* 1967), with some duplicated genes silenced after a lineage specific whole genome duplication (referred to here as “Cc4R”). Our aims in this study were to: (1) succinctly summarize and compare genes and functional annotation information obtained from various databases; (2) test whether *Cyprinus carpio* expresses additional copies of particular genes compared to the two diploid species *(Cyprinella lutrensis* and *Platygobio gracilis);* (3) identify potentially novel genes present in the three cyprinids but not found in zebrafish or other genome databases; and (4) to assess evolutionary conservation of essential genes for development.

In zebrafish, 307 genes are known to be essential for development. Knockout mutations in these genes are embryonic lethal according to experiments by Amsterdam *et al.* (2004), with subsequent revisions by Chen *et al.* (2012) and updates to the ENSEMBL database (Flicek *et al*. 2014). These genes are highly conserved across extremely deep phylogenetic branches (e.g., yeast, fly, zebrafish, and human) due to their essential roles in development (Amsterdam *et al*. 2004). Despite their importance, essential genes have not been studied in the context of comparative molecular ecology or ecological genomics of co-occurring species. Using transcriptome data presented in this study, we assessed the evolutionary conservation of the 307 zebrafish essential genes across three additional cyprinid lineages. We predicted that these genes would be highly conserved across all species, consistent with their critical functional roles. We also tested whether both copies of duplicated genes in *C. carpio* (i.e., Cc4R Ohnologs) were retained and expressed in duplicate or whether one copy was evolutionarily lost.

## MATERIALS AND METHODS

Fish (n = 3 per species) were collected with a seine on 6 July 2012 from a field site on the Rio Grande, approximately 40 km south of Socorro, New Mexico (33.690556°N, 106.993042°W). Whole fish samples were immediately frozen in liquid nitrogen and transported to the laboratory. Skin, gill, gut, and kidney tissues were dissected as they thawed, placed in TRIzol (Invitrogen), and mechanically homogenized. Total RNA was isolated using Purelink RNA Mini kits (Ambion) following manufacturer’s protocol, along with DNase treatment to reduce genomic DNA contamination. Purified total RNA was sent to the National Center for Genome Resources (Santa Fe, New Mexico, USA) for quantification, quality assessment, cDNA library preparation and sequencing. RNA integrity and purity was assessed with a Bioanalyzer 2100 instrument (Agilent Technologies). Thirty-six Illumina libraries were constructed (3 species x 4 tissues x 3 biological replicates) from the total RNA samples using Illumina TruSeq DNA prep kits according to the manufacturer’s protocol. Libraries were barcoded using standard six base pair Illumina oligonucleotides, and six libraries were pooled for each lane of Illumina HiSeq 2000 (V3 chemistry) for a total of six lanes of 2 x 100 bp paired-end sequencing.

### Bioinformatics

We developed a bioinformatics pipeline (Fig. 1) for analyzing transcriptomic data in three main steps: *de novo* assembly, gene annotation, and analysis of expression of duplicated genes. Adapters and barcode sequences were removed from raw reads, and reads were trimmed using TRIMMOMATIC (Bolger *et al.* 2014) with parameter settings as follows: leading quality = 5; trailing quality = 5; minimum trimmed read length = 36). Reads were normalized *in silico* to maximum read coverage of 50X. Clipped and trimmed reads were assembled, *de novo,* for each species separately using TRINITY (version 2014-04-13; Grabherr *et al.* 2011; Haas *et al.* 2013), with minimum contig length set to 200 bp. TRINITY assembles reads into contigs (“TRINITY transcripts”), places similar transcripts in groups loosely referred to as “genes”, and groups similar “genes” into gene clusters. *De novo* TRINITY assemblies were annotated using TRINOTATE, a comprehensive annotation package distributed with the TRINITY package suite. TRINOTATE was used to query contigs against the following databases or search tools: BLASTx, BLASTp, Pfam, SignalP, Uniprot, eggnog, and gene ontology.

**Figure 1.**
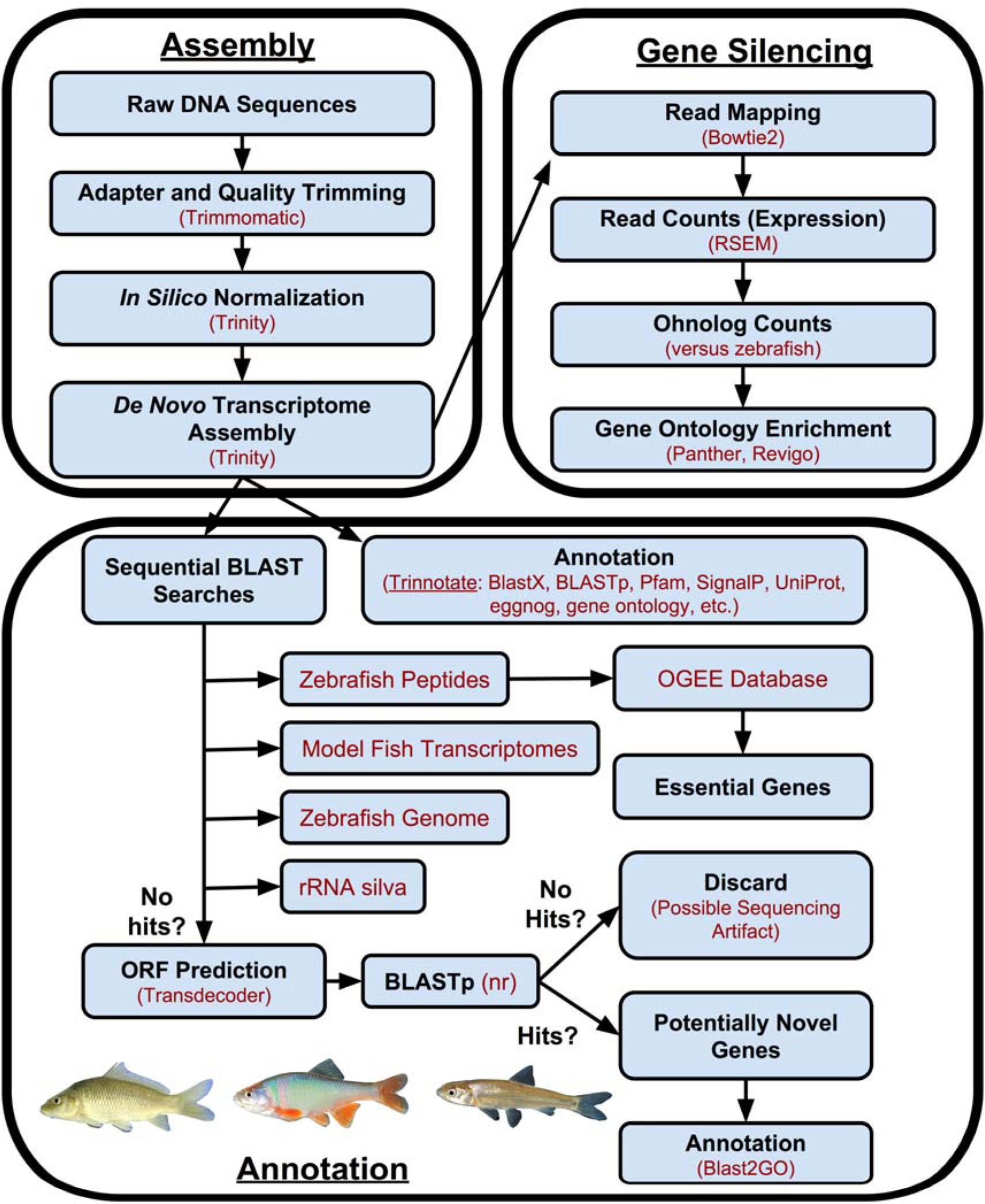
Flow diagram illustrating our bioinformatic pipeline. Analyses consist of three main steps: assembly, annotation, and analysis of gene silencing patterns. Databases queried and software packages used are in red font.

Putative protein coding genes were also identified by BLASTx searches of contigs against zebrafish *(Danio rerio)* peptide sequences (database build Zv9) obtained from Ensembl 78 (Flicek *et al.* 2014). Significant BLAST hits were identified based on the following parameter settings: E-value < 0.0001; gap open penalty = 11; gap extend = 1; wordsize = 3. After extensive testing, this parameter combination was found to give the optimal balance between finding matches for large numbers of contigs, while minimizing spurious hits. For most genes a 1-1 match was expected between zebrafish versus *Platygobio gracilis* or *Cyprinella lutrensis,* whereas zebrafish and *Cyprinus carpio* should have either 1-2 or 1-1 due to partial diploidy in carp. We used this expectation in deciding the threshold E-value to use. In practice, more stringent E-value thresholds (e.g., E < 1e-6) had very little effect on the number of significant BLAST hits.

Contigs with no significant BLAST hits against the zebrafish transcriptome were subjected to a series of stepwise BLASTn searches until significant hits were found (or not) in order to identify the possible sources of those sequences (e.g., microbiome) or to identify novel genes not present in the zebrafish genome. First, remaining contigs lacking significant hits against the zebrafish transcriptome were queried against the rRNA silva database (SSU Ref 119 NR99 and LSU Parc 119), which contains bacterial and eukaryotic rRNA sequences (Pruesse *et al.* 2007). Contigs with still no significant BLAST hits were then queried against a database containing all nine additional teleost fish transcriptomes (Amazon molly, *Poecilia formosa;* cavefish, *Astyanax mexicanus;* cod, *Gadus morhua*; fugu, *Takifugu rubripes*; medaka, *Oryzias latipes*; platyfish, *Xiphophorus maculates*; stickleback, *Gasterosteus aculeatus*; tetraodon, *Tetraodon nigroviridis;* tilapia, *Oreochromis niloticus)* from Ensembl 78. Contigs with no BLAST hits at this point were then BLASTed against the zebrafish genome (Zv9) using the “Top Level” sequences from Ensembl to identify possible genomic DNA contamination. Remaining contigs with no significant blast hits in any of these databases were piped to TRANSDECODER (Haas *et al.* 2013) to identify open reading frames (ORFs) that represent potentially novel genes. Default parameter settings were used with TRANSDECODER. The software generates predicted peptide sequences for contigs with ORFs. Predicted peptide sequences for the contigs with ORFs but no BLAST hits to the aforementioned databases were queried (BLASTp; e-value < 0.001) against the NCBI nr database. BLAST2GO (version 3.0, Conesa *et al.* 2005) was used to identify top species hits for those predicted proteins with significant hits against nr. The remaining sequences with no hits to databases and no ORFs were discarded as likely non-protein coding, genomic DNA contamination with sufficient evolutionary divergence from zebrafish to render genomic BLASTn searches ineffective.

### Genome duplication, diploidization and gene silencing

Trimmed sequence reads were mapped to TRINITY contigs using BOWTIE2 (version 2.2.2.3; Langmead & Salzberg 2012) and corresponding gene expression was quantified with RSEM (version 1.2.13; Li & Dewey 2011). Because RSEM is incompatible with indel, local, and discordant alignments, parameter settings were chosen to avoid these alignments. The following RSEM parameters were used: ‐‐sensitive; ‐‐dpad 0; ‐‐gbar 99999999; ‐‐mp 1,1 ‐‐np 1 ‐‐score-min L,0,-0.1; ‐‐no-mixed; ‐‐no-discordant. Normalized expression for TRINITY genes was calculated by standardizing by total mapped reads across libraries and summed across alternate TRINITY transcripts (isoforms) for each locus. In order to assess the expression of duplicated genes in *Cyprinus carpio* arising from the carp-specific whole genome duplication (“Cc4R”), we quantified the number of TRINITY genes present in each species relative to zebrafish genes, as well as their expression levels. We used an arbitrary threshold of ten sequence reads per gene per tissue, summed across all three individuals, for a given gene to be considered “expressed” in a particular tissue. This approach was aimed at reducing the influence of unique or nearly unique reads (e.g., sequencing artifacts). Most of the contigs excluded as a result of this filtering were contigs represented only by singleton reads in one library.

For *C. carpio,* we tested whether certain functional classes of genes were preferentially expressed in duplicate (i.e., the case where neither ohnolog silenced). For this analysis, we used PANTHER (MI *et al.* 2013) to test for statistical overrepresentation of GO-slim Biological Processes, with Bonferroni correction. The test genes consisted of the list of *C. carpio* ohnologs expressed in duplicate, while the list of all *C. carpio* genes present in the assembly was used as the reference set. GO terminology was based on the zebrafish database. Results of the overrepresentation analysis were visualized with REVIGO (Supek *et al.* 2011).

### Essential genes

To test the hypothesis of evolutionary conservation of essential genes among cyprinid fishes, we used zebrafish genes present in the Online Gene Essentiality Database (OGEE; Chen *et al.* 2012) and identified orthologs in the three transcriptomes from BLASTx searches described above. Of the 307 essential genes in zebrafish (Amsterdam *et al.* 2004; Chen *et al.* 2012), one (ENSDARG00000038423) has been retired from ENSEMBL and one (ENSDARG00000045605) is an unprocessed pseudogene with no protein product. We searched for the remaining 305 genes in the three transcriptome assemblies to assess their conservation across the cyprinid phylogeny. We predicted that expression would be conserved due to their high rate of conservation seen in other organisms (Amsterdam *et al.* 2004).

### Data accessibility

Raw sequence reads were uploaded to the NCBI Sequence Read Archive (Acc. Nos. SRXXXX, SRXXXX, SRXXXX). The three TRINITY assemblies were archived as separate bam files in NCBI Transcriptome Shotgun Assembly Database (Acc. Nos. XXXXX, XXXXX, XXXXX). Because these raw assemblies also contain microbiome and genomic sequences, we also deposited the final, filtered, high quality transcriptome fasta files in Dryad (Acc. Nos. XXXXX). These files include TRINITY contigs with significant BLASTx hits for zebrafish or the nine other fish transcriptomes, as well as companion fasta files containing the potentially novel genes (i.e., ORFs present but no significant BLAST hits in nr or the other databases). Annotations files from TRINOTATE were also deposited in Dryad.

## Results

### Sequencing and transcriptome assemblies

Six lanes of Illumina sequencing produced more than 1.2 billion paired-end reads, including 420.5-, 413.9-, and 385.3-million sequences in *Cyprinus carpio, Cyprinella lutrensis,* and *Platygobio gracilis,* respectively. *De novo* assembly resulted in high quality transcriptomes for all three species (Table 1). The *C. carpio* assembly had the largest number of contigs (“TRINITY transcripts”) and genes (“TRINITY genes”), while *P. gracilis* had the fewest. In contrast, metrics for contig length (N25, N50, N75, median contig length, average contig length) were all longer in *P. gracilis* than the other two species (Table 1). Overall, the *P. gracilis* transcriptome assembly was more complete despite fewer raw sequence reads. TRANSDECODER predicted ORFs in about half of all TRINITY contigs (Table 2), with the remainder comprised mainly of genomic DNA contamination that was filtered out of the final dataset. The N50 of predicted ORFs was only slightly shorter in the three species (i.e., 1,299 - 1,572 bp) than in zebrafish (CDS N50 = 2,037 bp), and similar to the recently published draft *C. carpio* genome (1,487 bp; Xu *et al.* 2014). Removal of microbiome and genomic DNA contamination from the final assembly resulted in fewer, but longer contigs (see filtering of the final dataset, below), and an overall higher-quality assembly.

**Table 1.** *De novo* transcriptome assembly results. Zebrafish *(Danio rerio)* data is included as an example of a well-assembled and complete transcriptome based primarily on Sanger sequencing.

|  | *Cyprinus*<br>*carpio* | *Cyprinella*<br>*lutrensis* | *Platygobio*<br>*gracilis* | *Danio*<br>*rerio* |
| --- | --- | --- | --- | --- |
| Trinity “genes”<br>(=Clusters of contigs) | 309,921 | 255,863 | 180,130 | 30,651 |
| Trinity “transcripts”<br>(=Assembled contigs) | 440,696 | 382,504 | 262,969 | 43,153 |
| GC content | 42.45 | 43.25 | 42.67 | 49.60 |
| N25 (bp) | 3,327 | 3,069 | 3,644 | 3,465 |
| N50 | 1,841 | 1,666 | 1,972 | 2,037 |
| N75 | 704 | 679 | 788 | 1,179 |
| Median contig length | 418 | 439 | 450 | 1,080 |
| Average contig length | 907 | 886 | 978 | 1,501 |
| Total assembled bases | 399,790,412 | 339,160,955 | 257,217,466 | 64,757,328 |

**Table 2.**
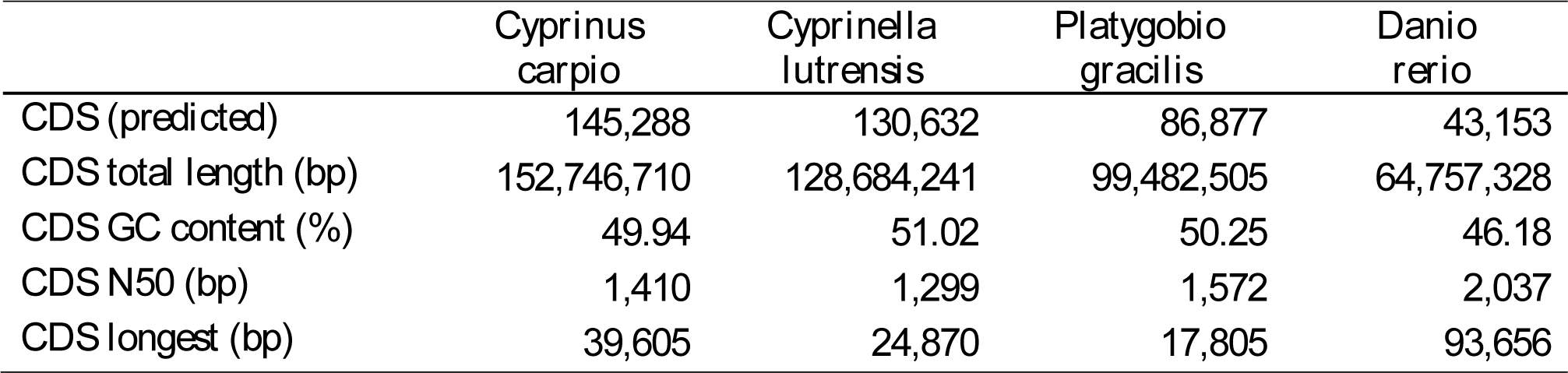
Assembly results and predicted coding sequence (CDS) in three study species and zebrafish. Zebrafish data is from ENSEMBL 78 (Flicek *et al.* 2014).

|  | *Cyprinus*<br>*carpio* | *Cyprinella*<br>*lutrensis* | *Platygobio*<br>*gracilis* | *Danio*<br>*rerio* |
| --- | --- | --- | --- | --- |
| CDS (predicted) | 145,288 | 130,632 | 86,877 | 43,153 |
| CDS total length (bp) | 152,746,710 | 128,684,241 | 99,482,505 | 64,757,328 |
| CDS GC content (%) | 49.94 | 51.02 | 50.25 | 46.18 |
| CDS N50 (bp) | 1,410 | 1,299 | 1,572 | 2,037 |
| CDS longest (bp) | 39,605 | 24,870 | 17,805 | 93,656 |

### BLAST searches: zebrafish transcriptome

Top BLASTx hits of TRINITY contigs against zebrafish peptides included approximately 20,000 unique genes (ENSDARG) and 11,000 protein families (ENSFAM) present in each of the three species (Fig. 3), suggesting similar annotation efficiency and transcriptome representation for each species. However, after pooling isoforms, the number of TRINITY genes that significantly matched these ~20,000 zebrafish genes varied among species: 66,447 in *Cyprinus carpio,* 60,990 in *Cyprinella lutrensis,* and 39,915 in *Platygobio gracilis* (Table 3, top row). Zebrafish genes were well covered, with more than 15,000 unique zebrafish genes covered over at least 70% of their length in corresponding contigs from each of the three cyprinids, consistent with the N50 data presented above. In general, zebrafish proteins were more completely covered by *P. gracilis* contigs than *C. carpio* or *C. lutrensis.* For example, zebrafish genes were more than 90% covered by sequences in 50.3% (12,489 of 24,817 genes) of *P. gracilis* genes with significant zebrafish peptide hits versus 49.9% (13,453 of 26,963) for *C. carpio* and 46.8% (12,538 of 26,817) in *C. lutrensis.* A large number of TRINITY contigs did not significantly match (BLASTx) zebrafish peptide sequences and were subsequently queried against several additional databases.

**Figure 3:**
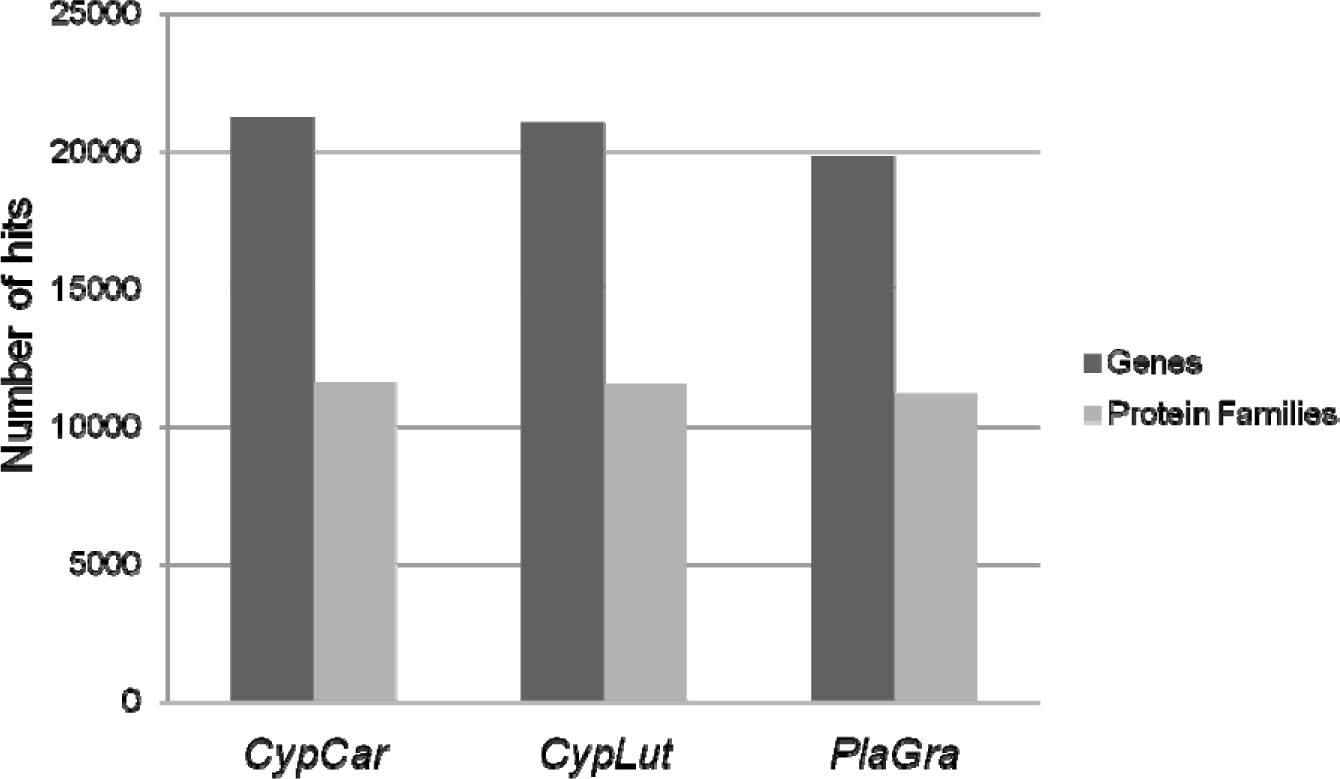
Unique genes and protein families from BLASTx searches (E-value threshold = 0.0001) against zebrafish *(Danio rerio)* peptide sequences. Similar numbers of zebrafish genes (~20,000) and protein families (~11,000) were identified across the three species, suggesting comparable assembly and annotation efficiency across these species.

**Table 3.** Significant BLAST hits for TRINITY “genes” versus zebrafish peptides, rRNA silva, (non-zebrafish) teleost fish transcriptomes, and the zebrafish genome. BLAST searches were done in stepwise fashion: all TRINITY genes were queried against zebrafish peptides but only genes without zebrafish peptide hits were queried against rRNA silva, and so on until all of the databases were queried.

|  | *Cyprinus*<br>*carpio* | *Cyprinella*<br>*lutrensis* | *Platygobio*<br>*gracilis* |
| --- | --- | --- | --- |
| Zebrafish peptides | 66,447 | 60,990 | 39,915 |
| rRNA Silva (microbiome) | 140 | 306 | 87 |
| Teleost fish transcriptomes | 4,572 | 2,923 | 1,561 |
| Zebrafish genome | 48,527 | 38,199 | 31,955 |
| No significant BLAST hits | 190,235 | 153,445 | 106,612 |
| **Total Contigs** | **309,921** | **255,863** | **180,130** |

### BLAST searches: other databases

Contigs lacking significant BLASTx hits against zebrafish peptides were queried (BLASTn) iteratively against rRNA silva microbiome database, nine teleost transcriptomes, and the zebrafish genome databases (Table 3). For contigs lacking hits against zebrafish peptides, BLASTn searches versus the rRNA silva database revealed a small number of significant hits (i.e., <400 contigs; Table 3). BLASTn searches of the remaining unmatched contigs versus the nine teleost fish transcriptomes identified approximately 1,500 - 4,500 additional hits (Table 3), far fewer than the evolutionarily more closely related zebrafish transcriptome. BLASTn searches of the remaining unidentified contigs against the zebrafish genome revealed a large number of significant hits (>30,000 per species), suggesting these reads were the result of low levels of background genomic DNA contamination in the cDNA libraries, a common occurrence resulting from the hypersensitivity of Illumina sequencing. Conservation of sequences across deep evolutionary lineages suggests functional importance, such as noncoding regulatory regions.

Despite extensive BLAST searches, a large number of TRINITY contigs (>100,000 in each species or more than 50% of all contigs) did not have significant hits in any of the databases. These, contigs are short in length (i.e., 200 bp) and have few reads mapping to them (e.g., single-read contigs). These could represent endogenous genomic DNA contamination of cDNA libraries and have sufficient evolutionary divergence from zebrafish to render BLASTn searches ineffective. A large number are also expected to be non-rRNA sequences from the microbiome, which were not present in target databases. Of contigs with no BLAST hits in the aforementioned databases, TRANSDECODER predicted ORFs in 8,652 *(Cyprinus carpio),* 9,215 *(Cyprinella lutrensis),* and 3,011 *(Platygobio gracilis)* contigs (Table 4). Roughly half of the predicted ORFs had significant BLASTp hits against the nr protein database (3,789, 4,154, and 1,548 contigs, respectively). Conversely, there were 4,863 *(C. carpio),* 5,061 *(C. lutrensis),* and 1,463 *(P. gracilis)* predicted ORFs had no significant hits against nr (Table 4). These ORFs could include novel genes not present in zebrafish or other teleost models, genes present in zebrafish but with significantly divergent sequences or exon structure to cause BLAST searches to miss them, or could include genes from the microbiome that are not present in sequence databases.

**Table 4.**
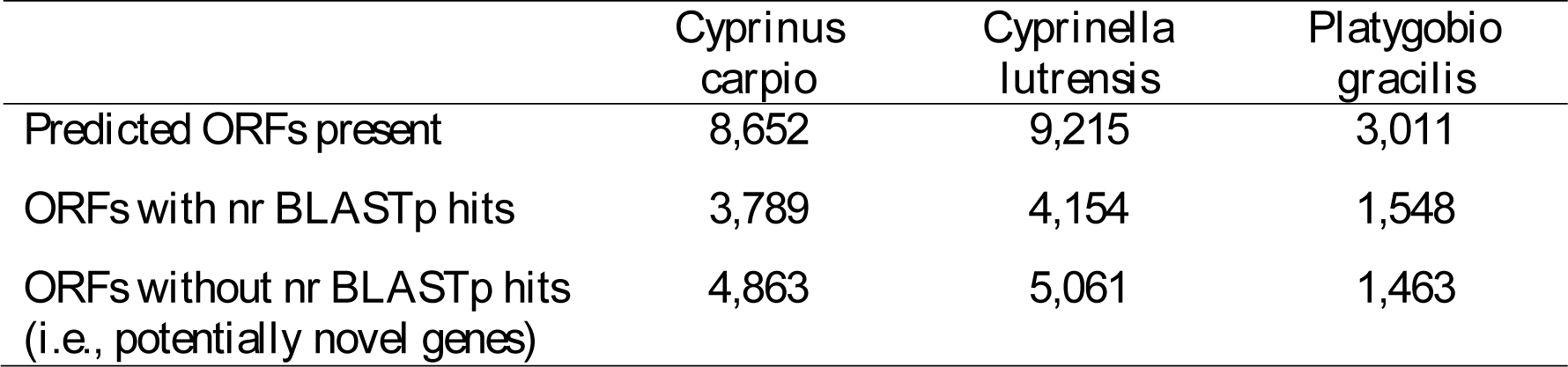
Summary of open reading frames (ORFs) identified in TRINITY contigs with no significant BLAST hits against databases listed in Table 3 (“No significant BLAST hits”). Roughly half of these predicted ORFs had significant protein level BLAST hits in the nr database. Some of the ORFs lacking similar proteins in the nr database may represent novel genes or genes with divergent sequences and function, while many are likely spurious results from the sequencing and assembly process or are from unidentified microbiota.

For ORFs with nr hits, zebrafish was the top-hit species for a large portion (Fig. 4), somewhat paradoxically given the lack of significant BLAST hits against zebrafish peptide and genome sequences discussed above. This appears to be due to the fact that TRANSDECODER-predicted ORFs exclude 5’ and 3’ untranslated regions (UTRs) which diverge more rapidly than ORFs over evolutionary time. In *C. carpio* and *C. lutrensis,* many of these ORFs are from the microbiome and share significant similarity to cyclophyllid tapeworms (e.g., *Echinococcus, Hymenolepis)* and protozoans (e.g., *Tetrahymena, Parameceum).* Conversely, in *P. gracilis* the ORFs appear to be endogenous genes with high similarity to zebrafish (Fig. 4). Contigs with predicted ORFs but no BLAST hits to any of the databases possibly represent novel or functionally divergent genes in these species that warrant further study. These sequences are available as fasta files in Dryad (XXXXXXXX).

**Figure 4.**
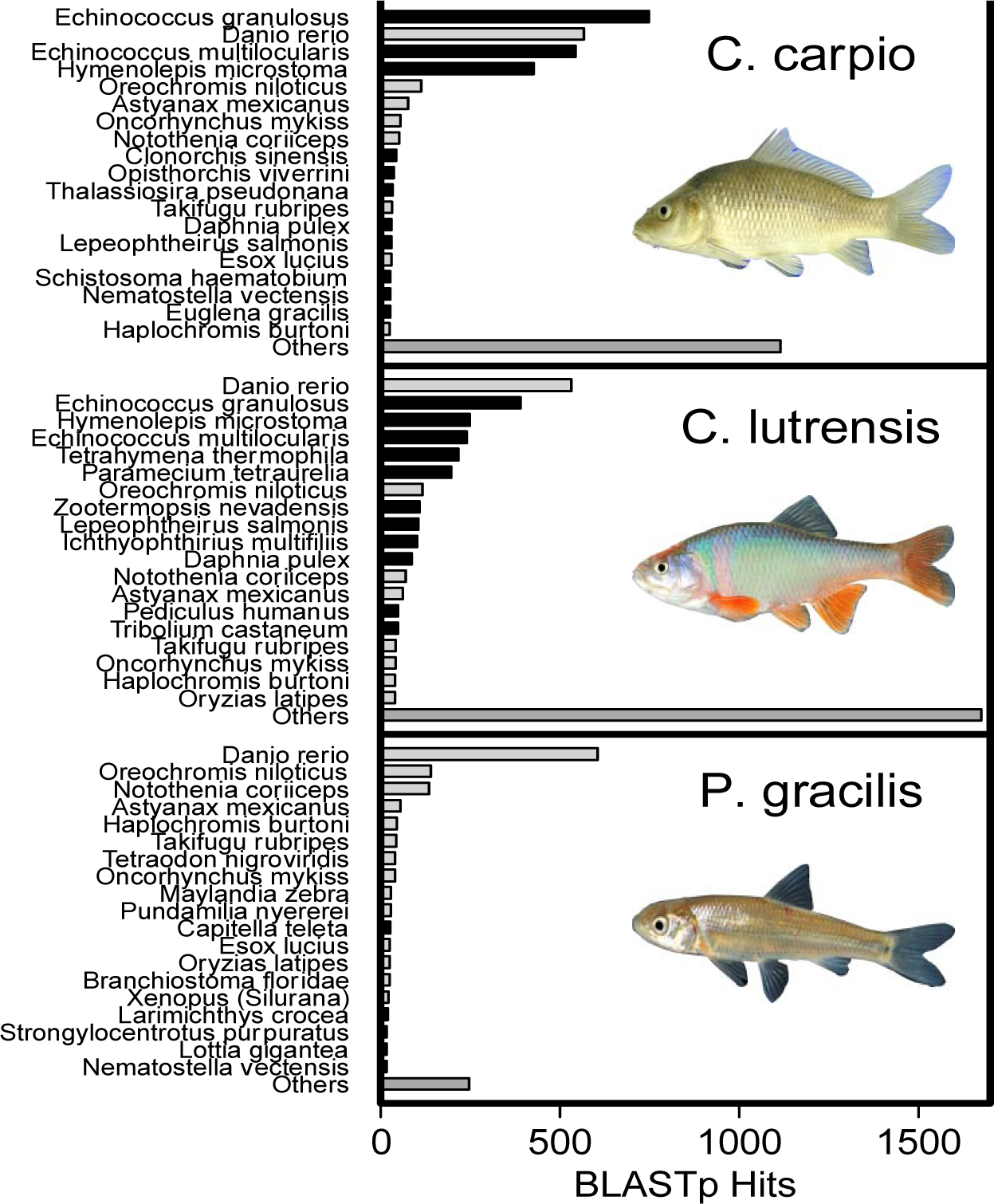
Top-species BLASTp hits for predicted open reading frame (ORF) peptide sequences queried against the nr database. Query sequences only included ORFs from contigs that lacked significant BLAST hits (see Tables 3 and 4). Grey bars represent fish or other chordates, while black bars represent non-chordate taxa. In *Cyprinus carpio* and *Cyprinella lutrensis,* many of these ORFs are likely from the microbiome (black bars) as they share significant similarity to cyclophyllid tapeworms (e.g., *Echinococcus, Hymenolepis)* and protozoans (e.g., *Tetrahymena, Parameceum).* Conversely, in *Platygobio gracilis* most of the ORFs appear to be endogenous genes with high similarity to zebrafish and other teleost fish.

### Filtering and the final assembly datasets

After filtering and removal of genomic DNA and microbiome reads, the final *de novo* assembly datasets contained only TRINITY contigs falling into one of the following categories: (1) contigs with significant BLAST hits against zebrafish or the nine other teleost transcriptomes; or (2) contigs with no matches against any of the databases but with predicted ORFs present, i.e., potentially novel genes. All other contigs were removed via bioinformatic filtering. The final datasets are significantly smaller than the raw *de novo* assembly deposited in NCBI TSA, but present much more reliable sequence information, i.e., actual transcriptome sequences rather than microbiome or genomic DNA contamination.

### Genome duplication, diploidization and gene silencing

Transcriptome annotation and comparison with zebrafish revealed that *Cyprinus carpio* expresses more genes than *Cyprinella lutrensis* and *Platygobio gracilis,* due to the carp-specific whole genome duplication (Fig. 5). *Cyprinus carpio* expressed about 41% more genes overall than *P. gracilis* and 11% more than *C. lutrensis.* The number of duplicate genes expressed varied dramatically among tissue types (Fig. 5). In all tissues except skin, *C. carpio* expressed more genes than the other two species (i.e., 3-48% more). In skin, both *C. lutrensis* and *P. gracilis* expressed more genes than *C. carpio* (26 and 2%, respectively). Using higher thresholds for “expression” had moderate impact on the inferred percentage of duplicates expressed: a threshold of 100 reads instead of 10 resulted in different estimates of duplicated genes expressed in *C. carpio* versus *P. gracilis* (18% more in *C. carpio)* and *C. lutrensis* (8% more in *C. carpio),* i.e., retained expression of Cc4R duplicates. The disparity in these results could be driven in part by different assembly qualities (e.g., a better assembled *P. gracilis* transcriptome).

**Figure 5.**
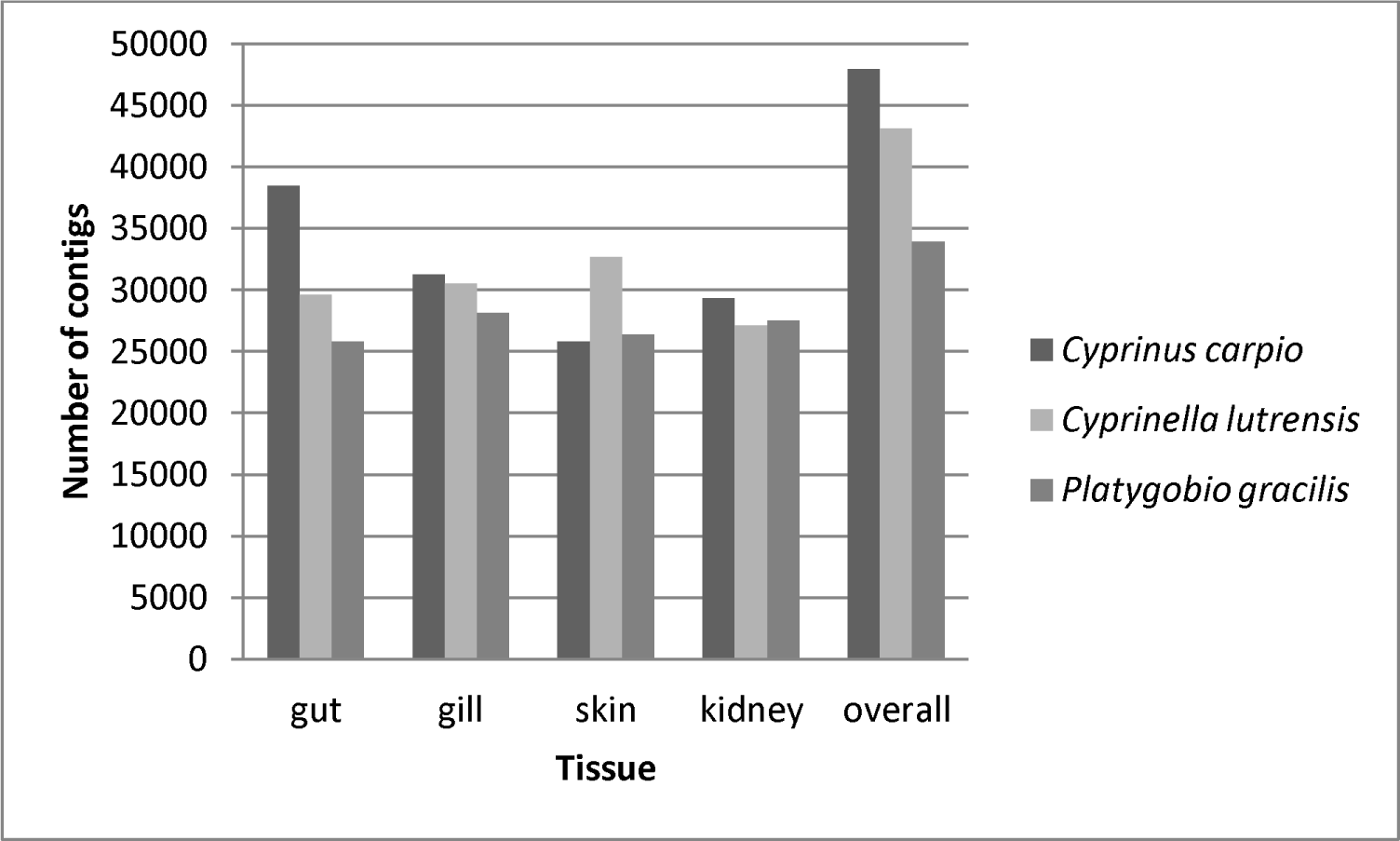
Number of TRINITY genes expressed in each of four tissue types, as well as all tissues pooled. Contigs only include those with significant BLASTx hits versus zebrafish peptides. Due to the carp-specific genome duplication event (Cc4R), *Cyprinus carpio* generally expresses more genes in a given tissue type than the other species, except in skin.

Genes with retained duplicate expression (i.e., Ohnologs) in *C. carpio* represented a diverse suite of functional groups: gene ontology terms that were significantly enriched in the ‘retained duplicates’ list were diverse (Fig. 6, top panel). One functional grouping that was a predominant contributor in the REVIGO analysis was “anatomical structure morphogenesis,” of interest because common carp attain much larger body size than the other two species (Fig. 6, bottom panel).

**Figure 6.**
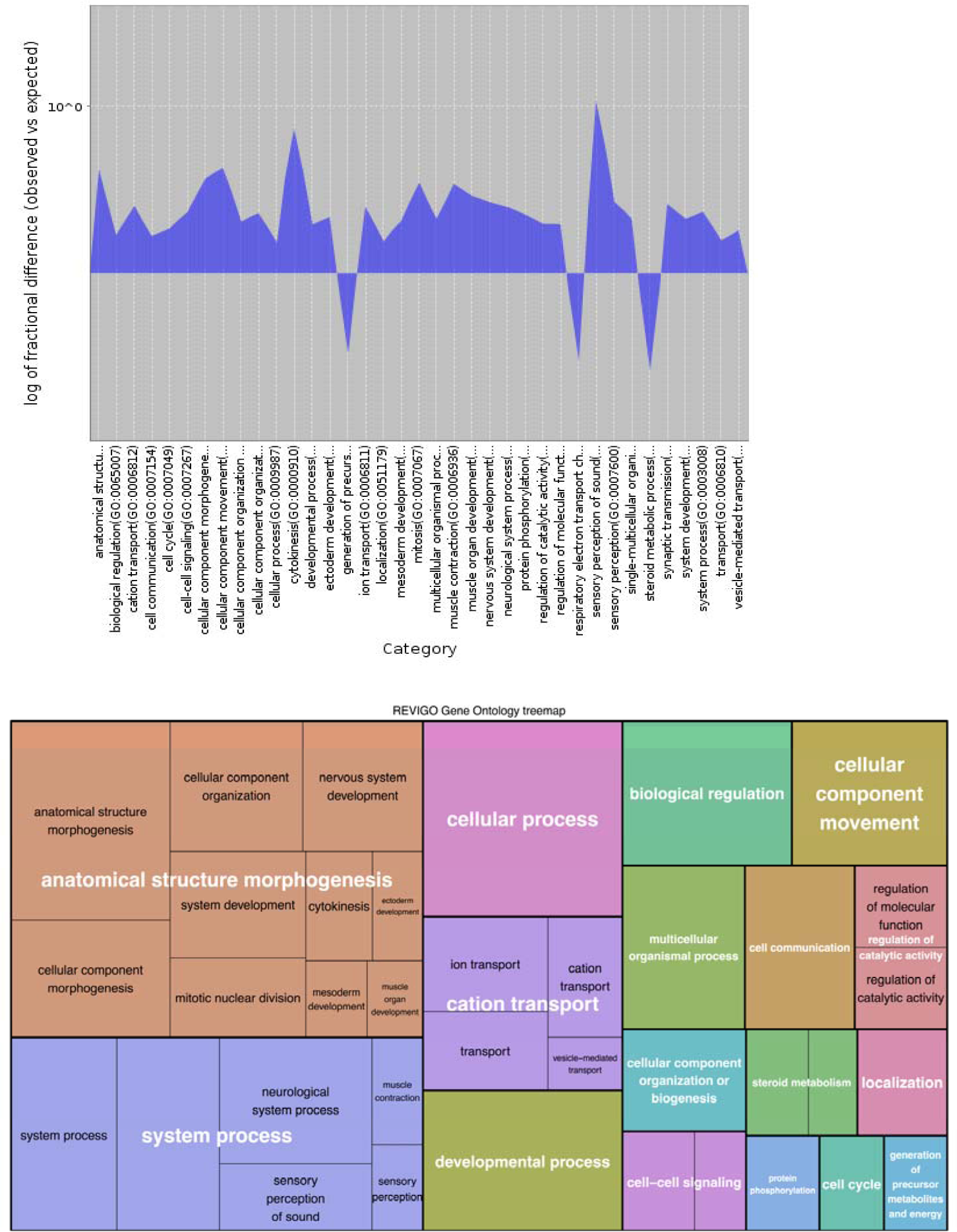
Top panel: Gene-ontology terms that are over‐ or under-represented (y-axis) in the list of genes retained as duplicates in the common carp transcriptome as compared to all expressed genes in common carp. Bottom panel: Summary of groups of biological processes overrepresented in the retained-duplicates in common carp. Box size is proportional to the number of genes with particular gene ontology terms, which may suggest a dosage effect in common carp.

### Expression of essential genes

Genes that are essential for embryonic development in *D. rerio* were nearly all present in the three cyprinids: 285 *(Platygobio gracilis),* 301 *(Cyprinella lutrensis),* and 301 *(Cyprinus carpio)* genes were expressed out of 305 zebrafish essential genes (i.e., 93.4 - 97.8%). Essential genes were nearly ubiquitously expressed across all four tissue types (skin, gill, gut, kidney), with low levels of tissue specificity (Fig. 7), in contrast to nonessential genes which generally exhibited higher levels of tissue specificity. Normalized levels of expression were higher in *C. carpio* than *P. gracilis* and *C. lutrensis* for 165 and 204 out of 305 genes, respectively. This pattern was not due to *C. carpio* expressing more loci per zebrafish gene (e.g., Ohnologs) than the other two species. Only slightly more loci (e.g., n=2 contigs) were expressed per essential gene in the recently duplicated *C. carpio* genome (Fig. 8) whereas most duplicated essential genes in *C. carpio* are not transcribed and have either been lost evolutionarily, e.g., pseudogenes, or are expressed in other developmental stages or tissues.

**Figure 7.**
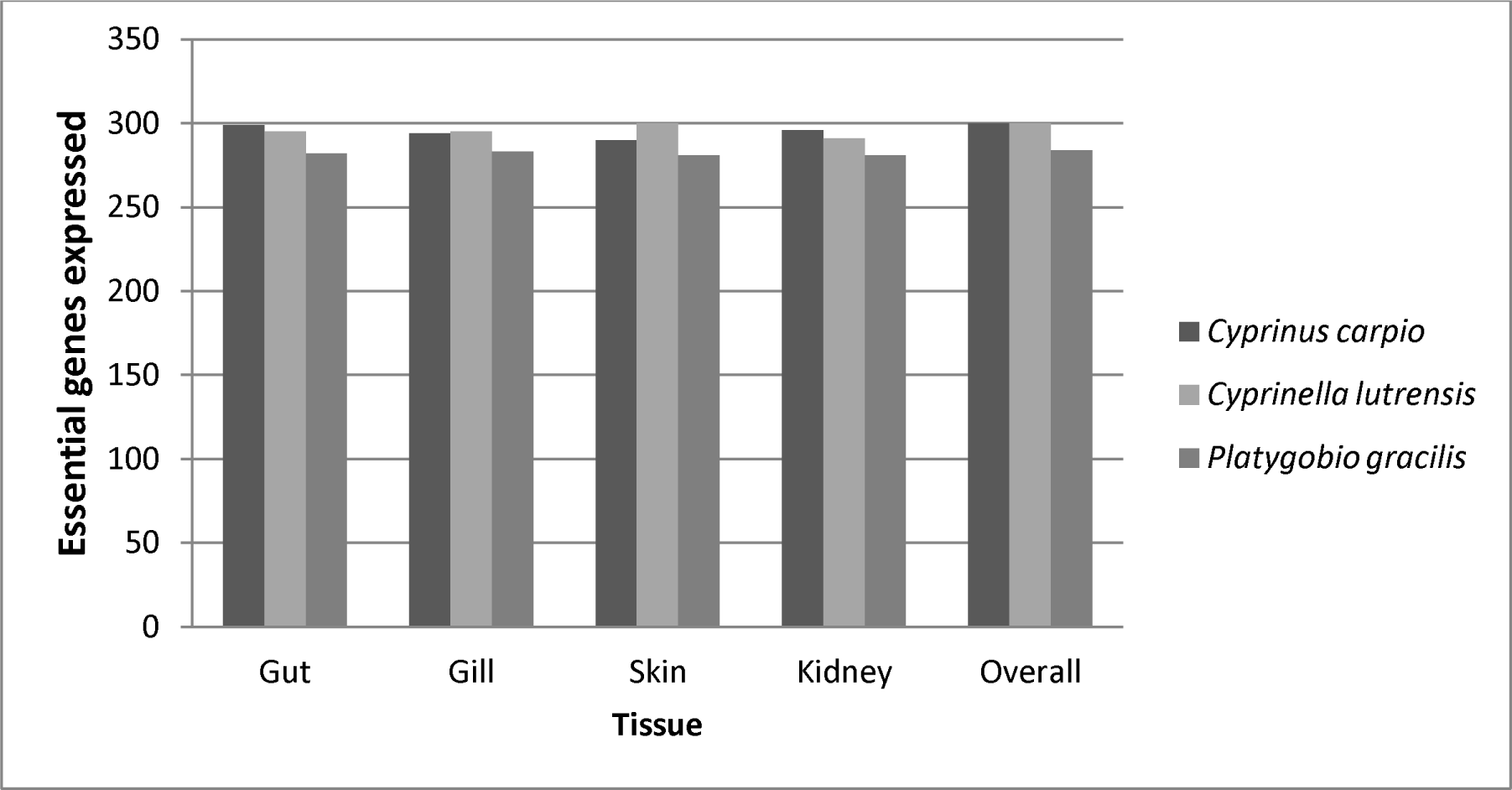
Expression of essential developmental genes by tissue type in three cyprinid fishes compared to 305 essential genes expressed across all tissues in zebrafish *(Danio rerio).* Essential genes were nearly ubiquitously expressed in all tissues.

**Figure 8.**
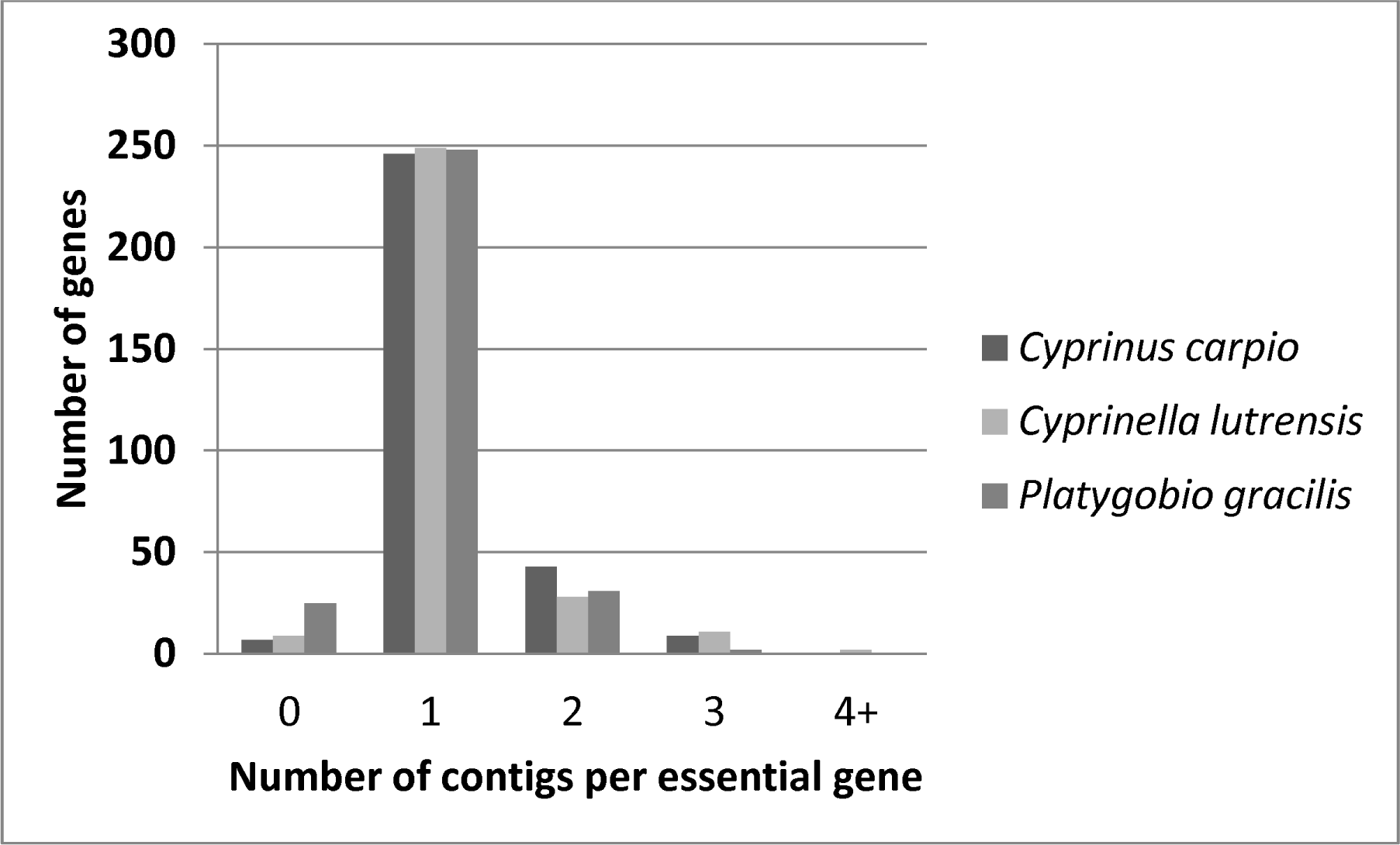
Number of loci (TRINITY genes) expressed per zebrafish essential gene. Only slightly more (e.g., n=2) were expressed per essential gene in the recently duplicated genome of *Cyprinus carpio.* Most duplicated essential genes in *C. carpio* are not transcribed and have either been lost evolutionarily, e.g., pseudogenes, or are expressed in other developmental stages or tissues.

## DISCUSSION

Next-generation transcriptome sequencing has revolutionized the field of molecular ecology over the past decade. One outcome is increased appreciation for the molecular complexity underlying the evolution of basic ecological traits. Here we demonstrated the utility and challenges associated with comparative study of transcriptomes in non-model organisms in a natural setting. Bioinformatic analyses requires careful processing and filtering to assess the sources of DNA fragments, which can be endogenous target transcriptome sequences, genomic DNA ‘contamination’ from the study organism, or DNA from the microbiome or diet items. Assessment of transcriptome quality also requires careful consideration. Traditional measures of assembled read lengths such as N50 are largely meaningless for transcriptomes without additional context. We advocate combining N50 and/or histograms of read lengths with explicit comparisons to well-studied transcriptomes of model organisms, when available. For example, we compared our *de novo* transcriptomes to zebrafish, which yielded valuable insight into progress made in our target species. Finally, positive identification of nearly all zebrafish essential genes in our transcriptomes is an additional test of our annotation procedures. Using the bioinformatics pipeline presented in Fig. 1, we demonstrate that high quality transcriptome data can be obtained from wild-caught organisms, which provide valuable tools for molecular ecologists studying functional genomics in a comparative context and natural settings.

We identified several key findings in this study, including: (1) high-quality transcriptome assemblies that reveal broad similarities and evolutionary conservation of genes with zebrafish, but with some key differences; (2) several potentially novel genes not identified in zebrafish that are candidates for studies of ecological novelty; (3) diverse microbiomes that vary strongly among the three species, despite their origin from a single collection locality; (4) extreme conservation of expression of essential genes for development; (5) a large number of duplicate genes expressed in the tetraploid, *Cyprinus carpio,* representing a diverse suite of biological processes or gene ontologies. We discuss each of these findings in greater detail below.

### Assembly results

There are important considerations associated with conducting transcriptome analysis in a non-laboratory setting and in species lacking high-quality, well-annotated genomes. For example, it is necessary to identify ways to maximize the quality and completeness of *de novo* assemblies. Our assemblies are somewhat less complete than the zebrafish reference, as expected because zebrafish has been sequenced extensively at the genomic DNA level, empirically validated with RNA-seq, and refined by years of manual curation.

TRINITY assemblies resulted in proportionally fewer long contigs (e.g., > 1000 bp) compared to zebrafish. Four factors account for this result. First, the microbiome is present in these sequences and many of the contigs are not endogenous, as reflected by top species hits in BLAST searches (Fig. 4). Second, a small amount of genomic DNA contamination persists despite DNase treatment during library preparation. Genomic contamination tends to be observed as short (e.g., 200 bp), shallow contigs often comprised of single-reads. Third, the *de novo* assemblies are more fragmented due to the short read technology employed, with multiple contigs often representing non-overlapping fragments of the same gene. This effect is particularly acute in genes with short sequence repeats, such as microsatellites. Finally, we only used the canonical zebrafish transcripts in this study, which excludes the shorter isoforms present in many genes and biases the zebrafish distribution toward longer sequences. Transcriptomes presented here represent an improvement (i.e., more sequences, higher coverage; longer relative N50) over earlier work on sequencing and assembling the common carp transcriptome using Roche 454 sequencing (Ji *et al.* 2012; Wang *et al.* 2012), due to the higher throughput, Illumina paired-end sequencing approach we employed. The bioinformatic approach we presented to identify and filter non-target sequences from the final dataset resulted in high quality and well annotated assemblies.

### Potentially novel genes

Results of BLAST searches and ORF predictions helped us identify candidate genes that may represent novel species‐ or taxon-specific genes. Our interest in these genes lies in the idea that they may contain some of the functional elements responsible for extensive ecological and phylogenetic diversity present in Cyprinidae. Many candidates may prove to be false positives as more fish genomes are sequenced and annotated; however, these candidates would be an excellent starting point for researchers interested in targeted searches for genes or proteins underlying ecological novelty in cyprinids that may have arisen through local gene duplications, exon shuffling, horizontal transfer, or other mechanisms.

### Microbiome diversity

Another valuable aspect of transcriptome sequencing of samples taken from nature is the simultaneous generation of quantifiable data on the microbiome. These data are applicable to study of host-parasite dynamics, immune response, paired comparative population genetics or phylogeographic analysis of host and microbiota. When generating *de novo* transcriptome assemblies for focal species, it is imperative that microbiome sequences are identified and filtered out of final assemblies. Genome-scale sequence data is often lacking for the bacterial and metazoan microbiota on vertebrate samples, which complicates attempts at removal. We used an iterative and successive filtering approach to address this issue (Fig. 1) that provides valuable information on the likely source (e.g., exogenous or endogenous) of particular sequences or contigs. Transcriptome characterization studies often do not attempt to remove exogenous microbiome and genomic DNA contamination. Researchers should be cautious when using unfiltered sequence reads, particularly when they are compiled into massive databases that lack appropriate metadata.

### Conservation of essential genes

Genes that are essential for embryonic development present interesting targets for studying genome evolution due to their critical functional importance. Zebrafish essential genes in the Online Gene Essentiality Database (OGEE; Chen *et al.* 2012) were widely transcribed in all three cyprinids and with low levels of tissue specificity. Coverage of essential genes is a metric that should be used to assess the quality and coverage of *de novo* transcriptome assemblies, analogous to the use of “housekeeping” genes as positive controls in qPCR studies. The presence of nearly all essential genes across these four cyprinid species (representing more than 100 million years of evolutionary time; Saitoh *et al.* 2011) is consistent with the hypothesis of broad evolutionary and functional conservation. The few essential genes not detected may still be present, but expressed transiently at larval or juvenile developmental stages. We propose that the number of essential genes expressed could be used as a metric to complement other measures of assembly quality and completeness, such as comparing transcript length histograms to closely related model species (see Fig. 2).

**Figure 2.**
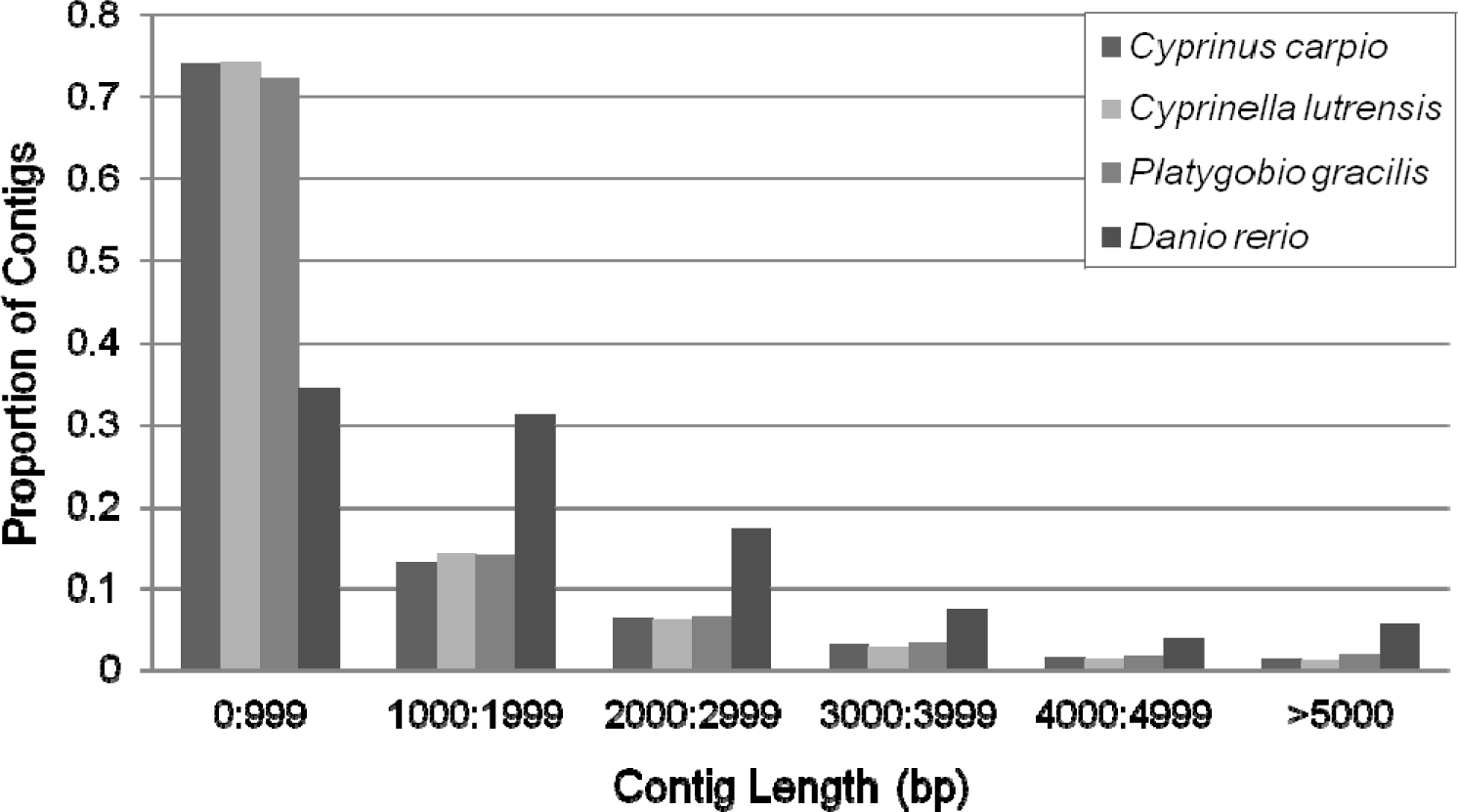
Contig length histogram of three cyprinids in this study and zebrafish, *Danio rerio.* By leveraging high throughput sequencing and bioinformatic filtering, we were able to generate high quality transcriptomes at a fraction of the cost and research effort used for zebrafish. As expected, *de novo* TRINITY assemblies resulted in proportionally fewer contigs longer than 1000 bp, as compared to those of a well-assembled transcriptome, zebrafish *(Danio rerio).* However, note that we only used canonical transcripts for zebrafish and not the shorter isoforms, which skews the distribution toward longer transcripts for that species.

### Tetraploidy and expression of duplicated genes

Our analyses suggested that a large number of duplicate genes are expressed in *Cyprinus carpio,* representing a diverse suite of biological processes or gene ontologies. Note that this analysis is based only on expressed genes in these tissues at a particular time point, rather than genomic DNA sequences and consequently would not include Ohnologs expressed only in different tissues or at different time points. Short reads in overlapping Ohnologous regions may not contain sufficient sequence divergence in recently duplicated genomes such as common carp, resulting in assembly of non-orthologous reads in the *de novo* assembly. The relatively more fragmented transcriptome of *Cyprinus carpio* in this study may reflect the challenge of assembling a recently duplicated transcriptome with little time for divergence in Ohnologs (e.g, 8.2 million years, Xu *et al.* 2014). In salmonids, although the fourth round of whole genome duplication is much older (i.e., 90-102 ma; Berthelot *et al.* 2014); many Ohnologous loci are still difficult to separate and some even maintain tetrasomic inheritance (Timusk *et al.* 2011). The draft genome sequence for common carp (Xu *et al.* 2014) should help identify Ohnologs that are in fact silenced by identifying pseudogenes in genomic DNA sequence. Previously, Wang *et al.* (2012) identified enrichment of retained duplicates in gene ontology pathways involved in immune function. Conversely, our analysis suggests for genes where both Ohnologs were expressed in *C. carpio,* there was enrichment in several different functional pathways, but in particular many genes were associated with “anatomical structure morphogenesis” (Fig. 6). This difference may be due to the large body size and rapid growth in *C. carpio* as compared to *C. lutrensis* and *P. gracilis* and an associated dosage effect. Ultimately, knowledge of which genes are retained and expressed in duplicate in tetraploids as compared to related diploid species can provide insight into the role that whole genome duplication plays in the molecular ecology and phylogenetic diversification of organisms.

## Summary

We present a bioinformatic analysis of short read sequences that yields high-quality transcriptome assemblies for non-model, ecologically-relevant organisms in a natural setting. The bioinformatics solutions presented here refine the process of gene annotation, assembly quality assessment, orthology assignment, and identification and partitioning of exogenous DNA in a single informatics pipeline. This approach facilitates technology transfer from a model organism (zebrafish) to a group of related species that fill diverse and important roles in these ecosystems and comprise an important component of biodiversity. The conserved expression of essential developmental genes across a broad phylogenetic scope and array of tissue types, and illustrated their utility as benchmarks for assessing coverage in *de novo* assemblies. More broadly, we demonstrate the promise and challenges associated with adapting model organism data to non-model, wild-caught samples.

## Acknowledgements

This project was supported by the National Institute of General Medical Sciences (8P20GM103451-12), New Mexico IDeA Networks of Biomedical Research Excellence (NMINBRE_A2_Jan_2013), and the Center for Evolutionary and Theoretical Immunology. Samples were collected under New Mexico Department of Game and Fish permit #3015. This research was approved by Institutional Animal Care and Use Committee Protocol #10-100468-MCC and #10-100492-MCC. We thank E. Loker, R. Miller, J. Kavka, and G. Rosenberg for research support. Thanks to Z. Ren and L. Hao for assistance with the Database of Essential Genes. Fish images were provided T. Kennedy (red shiner, flathead chub) and C. Thomas (common carp). This research benefitted from insight and technical assistance provided by F. Schilkey, N. Devitt, P. Mena, T. Ramaraj, I. Lindquist, A. Snyder, M. Osborne, and C. Krabbenhoft.

## Data Accessibility

Raw DNA Sequences: NCBI SRA (SRXXXX, SRXXXX, and SRXXXX). Transcriptome assemblies: NCBI TSA (XXXXXXX, XXXXXXX, and XXXXXXX). Final, filtered assemblies, fasta files of contigs representing to potentially novel genes, and TRINOTATE annotation output files are available from Dryad (Acc. Nos. XXXXXX, XXXXXX, and XXXXXX).

## Author Contributions

TJK and TFT designed and performed the research, TJK conducted the analyses, and TJK and TFT wrote the manuscript.

## REFERENCES

Amsterdam A, Nissen RM, Sun Z, et al.(2004) Identification of 315 genes essential for early zebrafish development. Proc Natl Acad Sci USA101, 12792–12797.

Berthelot C, Brunet F, Chalopin D, et al.(2014) The rainbow trout genome provides novel insights into evolution after whole-genome duplication in vertebrates. Nat Commun5, 3657.

Bolger AM, Lohse M, Usadel B (2014) Trimmomatic: a flexible trimmer for Illumina sequence data. Bioinformatics,btu170.

Briggs JP (2002) The zebrafish: a new model organism for integrative physiology. American Journal of Physiology-Regulatory, Integrative and Comparative Physiology282, R3–R9.

Chen W-H, Minguez P, Lercher MJ, Bork P (2012) OGEE: an online gene essentiality database. Nucleic Acids Res40, D901–D906.

Conesa A, Gotz S, Garcia-Gomez JM, et al.(2005) Blast2GO: a universal tool for annotation, visualization and analysis in functional genomics research. Bioinformatics21, 3674–3676.

Dooley K, Zon LI (2000) Zebrafish: a model system for the study of human disease. Curr Opin Genet Dev 10, 252–256.

Flicek P, Amode MR, Barrell D, et al.(2014) Ensembl 2014. Nucleic Acids Res42, D749–755.

Grabherr MG, Haas BJ, Yassour M, et al.(2011) Full-length transcriptome assembly from RNA-Seq data without a reference genome. Nat Biotechnol29, 644–652.

Haas BJ, Papanicolaou A, Yassour M, et al.(2013) De novo transcript sequence reconstruction from RNA-seq using the TRINITY platform for reference generation and analysis. Nature protocols8, 1494-1512.

Howe K, Clark MD, Torroja CF, et al.(2013) The zebrafish reference genome sequence and its relationship to the human genome. Nature496, 498–503.

Ji P, Liu G, Xu J, et al.(2012) Characterization of common carp transcriptome: *de novo*sequencing, assembly, annotation and comparative genomics. PLoS One7, 1–9.

Langmead B, Salzberg SL (2012) Fast gapped-read alignment with Bowtie 2. Nat Methods9, 357–359.

Li B, Dewey CN (2011) RSEM: accurate transcript quantification from RNA-Seq data with or without a reference genome. BMC Bioinformatics12, 323.

Lieschke GJ, Currie PD (2007) Animal models of human disease: zebrafish swim into view. Nature Reviews Genetics8, 353–367.

Mi H, Muruganujan A, Thomas PD (2013) PANTHER in 2013: modeling the evolution of gene function, and other gene attributes, in the context of phylogenetic trees. Nucleic Acids Res41, D377–386.

Ohno S, Muramoto J, Christian L, Atkin NB (1967) Diploid-tetraploid relationship among old-world members of the fish family Cyprinidae. Chromosoma23, 1–9.

Pruesse E, Quast C, Knittel K, et al.(2007) SILVA: a comprehensive online resource for quality checked and aligned ribosomal RNA sequence data compatible with ARB. Nucleic Acids Res35, 7188-7196.

Saitoh K, Sado T, Doosey MH, et al.(2011) Evidence from mitochondrial genomics supports the lower Mesozoic of South Asia as the time and place of basal divergence of cypriniform fishes (Actinopterygii: Ostariophysi). Zoological Journal of the Linnean Society161, 633–662.

Supek F, Bosnjak M, Skunca N, Smuc T (2011) REVIGO summarizes and visualizes long lists of gene ontology terms. PLoS One6, e21800.

Timusk ER, Ferguson MM, Moghadam HK, et al.(2011) Genome evolution in the fish family salmonidae: generation of a brook charr genetic map and comparisons among charrs (Arctic charr and brook charr) with rainbow trout. BMC Genet12, 1–15.

Wang J-T, Li J-T, Zhang X-F, Sun X-W (2012) Transcriptome analysis reveals the time of the fourth round of genome duplication in common carp (*Cyprinus carpio)*. BMC Genomics13, 96.

Xu P, Zhang X, Wang X, et al.(2014) Genome sequence and genetic diversity of the common carp, Cyprinus carpio. Nat Genet46, 1212–1219.

